# Genome-wide association and HLA region fine-mapping studies identify susceptibility loci for multiple common infections

**DOI:** 10.1101/073056

**Authors:** Chao Tian, Bethann S. Hromatka, Amy K Kiefer, Nicholas Eriksson, Joyce Y Tung, David A. Hinds

**Author notes:** Corresponding authors Email address: CT.

## Abstract

We performed 23 genome-wide association studies for common infections, including chickenpox, shingles, cold sores, mononucleosis, mumps, hepatitis B, plantar warts, positive tuberculosis test results, strep throat, scarlet fever, pneumonia, bacterial meningitis, yeast infections, urinary tract infections, tonsillectomy, childhood ear infections, myringotomy, measles, hepatitis A, rheumatic fever, common colds, rubella and chronic sinus infection, in more than 200,000 individuals of European ancestry. For the first time, genome-wide significant associations (*P* < 5 × 10^−8^) were identified for many common infections. The associations were mapped to genes with key roles in acquired and innate immunity *(HLA, IFNA21, FUT2, ST3GAL4, ABO, IFNL4, LCE3E, DSG1, LTBR, MTMR3, TNFRSF13B, TNFSF13B, NFKB1, CD40*) and in regulation of embryonic developmental process *(TBX1, FGF, FOXA1 and FOXN1).* Several missense mutations were also identified (in *LCE5A, DSG1, FUT2, TBX1, CDHR3, PLG, TNFRSF13B, FOXA1, SH2B3, ST5* and *FOXN1*). Missense mutations in *FUT2* and *TBX1* were implicated in multiple infections. We applied fine-mapping analysis to dissect associations in the human leukocyte antigen region, which suggested important roles of specific amino acid polymorphisms in the antigen-binding clefts. Our findings provide an important step toward dissecting the host genetic architecture of response to common infections.

## Introduction

Infectious diseases, the second leading cause of death worldwide, represent persistent challenges to human health due to increasing resistance to established treatments, lack of life-saving vaccines and medications in developing countries, and increasing distribution^1^. Studies have linked susceptibility to infectious agents to cancers, autoimmune diseases, and drug hypersensitivity. Human papillomaviruses are associated with multiple cancers^2^; rubella and mumps infection have been linked to development of type 1 diabetes (T1D) in children^3^; and risk of developing multiple sclerosis (MS) was greater among patients with herpes zoster (shingles) than in matched controls^4^. Reactivation of chronic persistent human herpesviruses has been linked to drug-induced hypersensitivity^5^. Thus, infectious diseases have a profound impact on our health, both directly as well as through connections with other diseases. Nevertheless, genetic studies of common infectious diseases lag somewhat behind those of other major complex diseases. Few genome-wide association studies (GWAS) have been undertaken for common infectious diseases of lower mortality or for infectious diseases for which vaccines are available^6^.

In this study, we conducted 23 imputed GWASs with more than 200,000 European research participants who were genotyped on the 23andMe platform and were asked to report on their history of infections (Table 1 and Supplementary note). Classical human leukocyte antigen (HLA) loci have been prototypical candidate genetic susceptibility to multiple infectious diseases^7^. We further imputed and tested the HLA classical alleles and amino acid polymorphisms to dissect independent HLA signals for multiple common infections that have significant HLA associations in our GWASs.

**Table 1.** Discovery cohort characteristics

| Phenotypes with significant GWAS findings |  |  |  |  |  |  |
| --- | --- | --- | --- | --- | --- | --- |
|  | Chicken pox | Shingles | Cold sores | Mononucleosis | Mumps | Hepatitis B |
| Cases | 107,769 | 16,711 | 25,108 | 17,457 | 31,227 | 1,425 |
| Control | 15,982 | 118,152 | 63,332 | 68,446 | 54,153 | 218,180 |
|  | Plantar warts | Positive TB | Strep throat<sup>a</sup> | Scarlet fever | Pneumonia | Bacterial meningitis |
| Cases | 24,994 | 4,426 | [qt] 52,487 | 6,812 | 40,600 | 842 |
| Control | 37,451 | 84,290 | 22,017 | 113,837 | 90,039 | 82,778 |
|  | Yeast infections<sup>a</sup> | UTI frequency<sup>a</sup> | Tonsillectomy | Childhood ear infection | Myringotomy |  |
| Cases | [qt] 52,218 | [qt] 35,000 | 60,098 | 46,936 | 4,138 |  |
| Control | 10,235 | 33,478 | 113,323 | 74,874 | 85,089 |  |
| Phenotypes with no significant GWAS findings |  |  |  |  |  |  |
|  | Measles | Hepatitis A | Rheumatic fever | # of Colds last year<sup>a</sup> | Rubella | Chronic sinus infection |
| Cases | 38,219 | 2,442 | 1,115 | [qt] 43,826 | 12,000 | 5,291 |
| Control | 47,279 | 217,137 | 88,076 | 15,720 | 71,597 | 79,622 |

<sup>a</sup>. Strep throat, yeast infections, urinary tract infection (UTI) frequency and number of colds last year were quantitative traits ([qt]).

## Results

In total, 59 genome-wide significant (GWS) regions (P < 5 × 10^−8^) were discovered (Table 2, see also Supplementary Figure 1 and Supplementary Figure 2 for GWAS Manhattan plots and regional plots). The HLA region is significantly associated with 13 of the infections that we studied and we dissected independent signals in each association (Figure 1 and Table 3, see also Supplementary Figure 3 for HLA regional plots). In the later sections, we describe the result for each phenotype in detail.

**Table 2:**
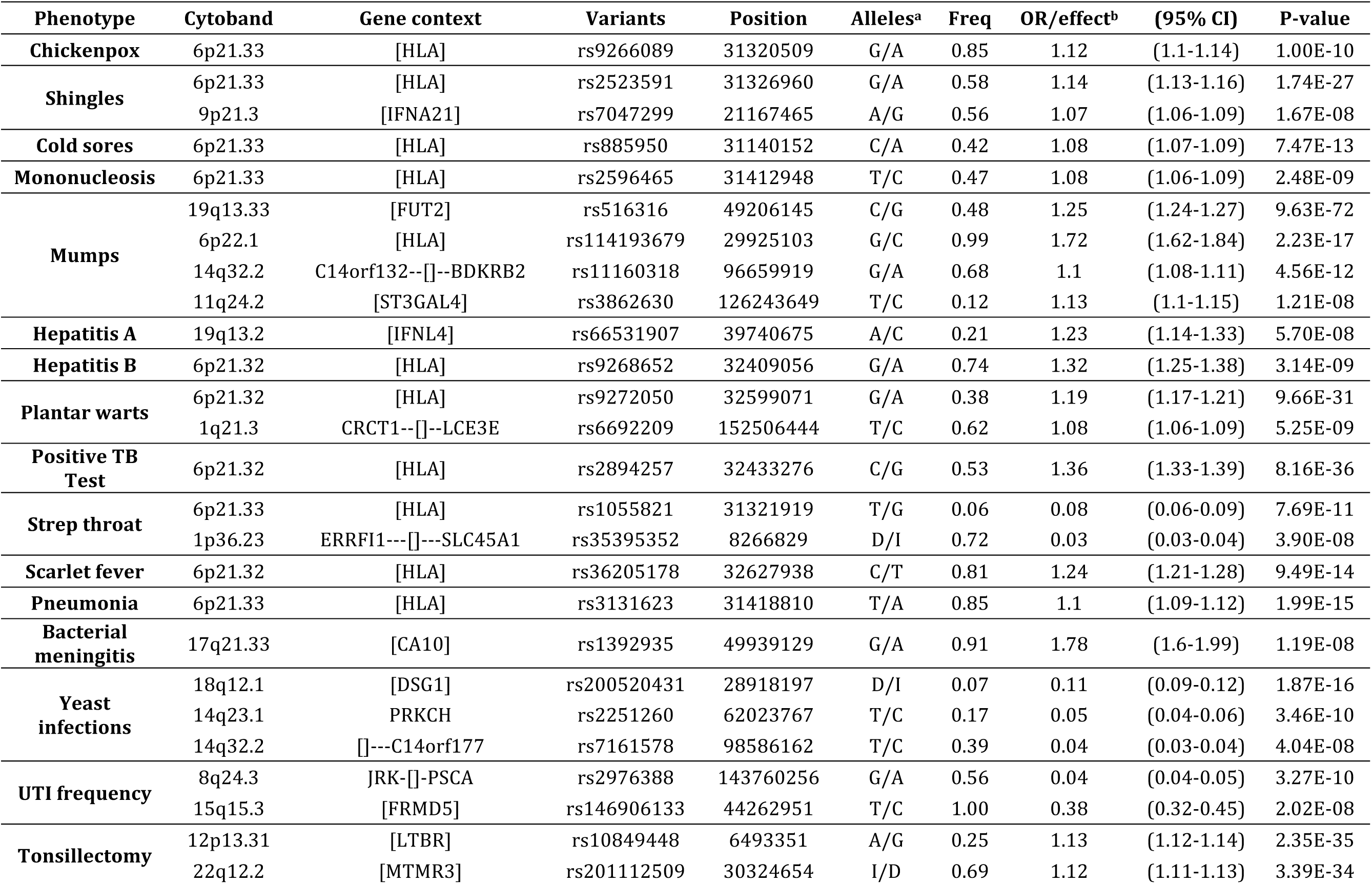
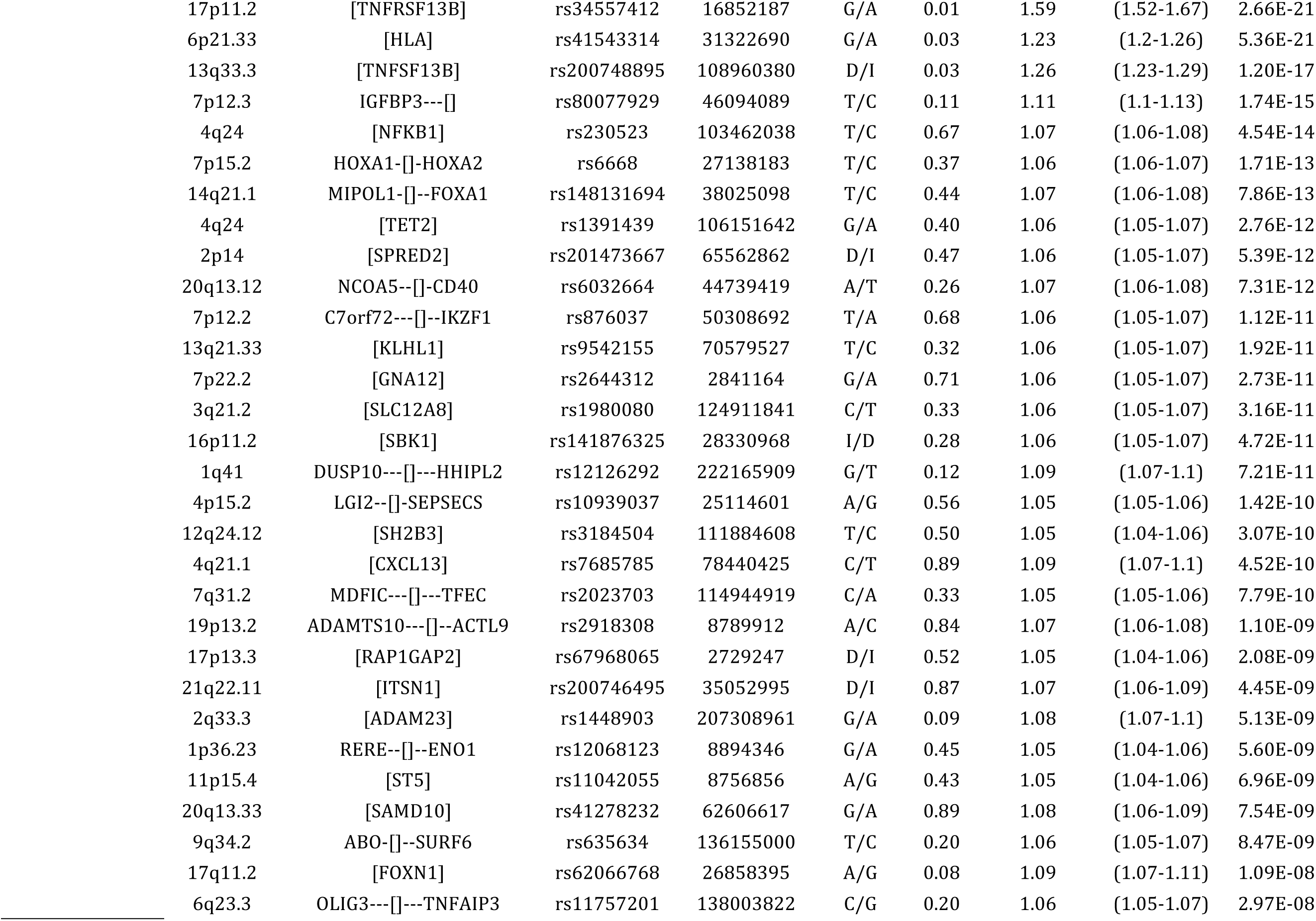
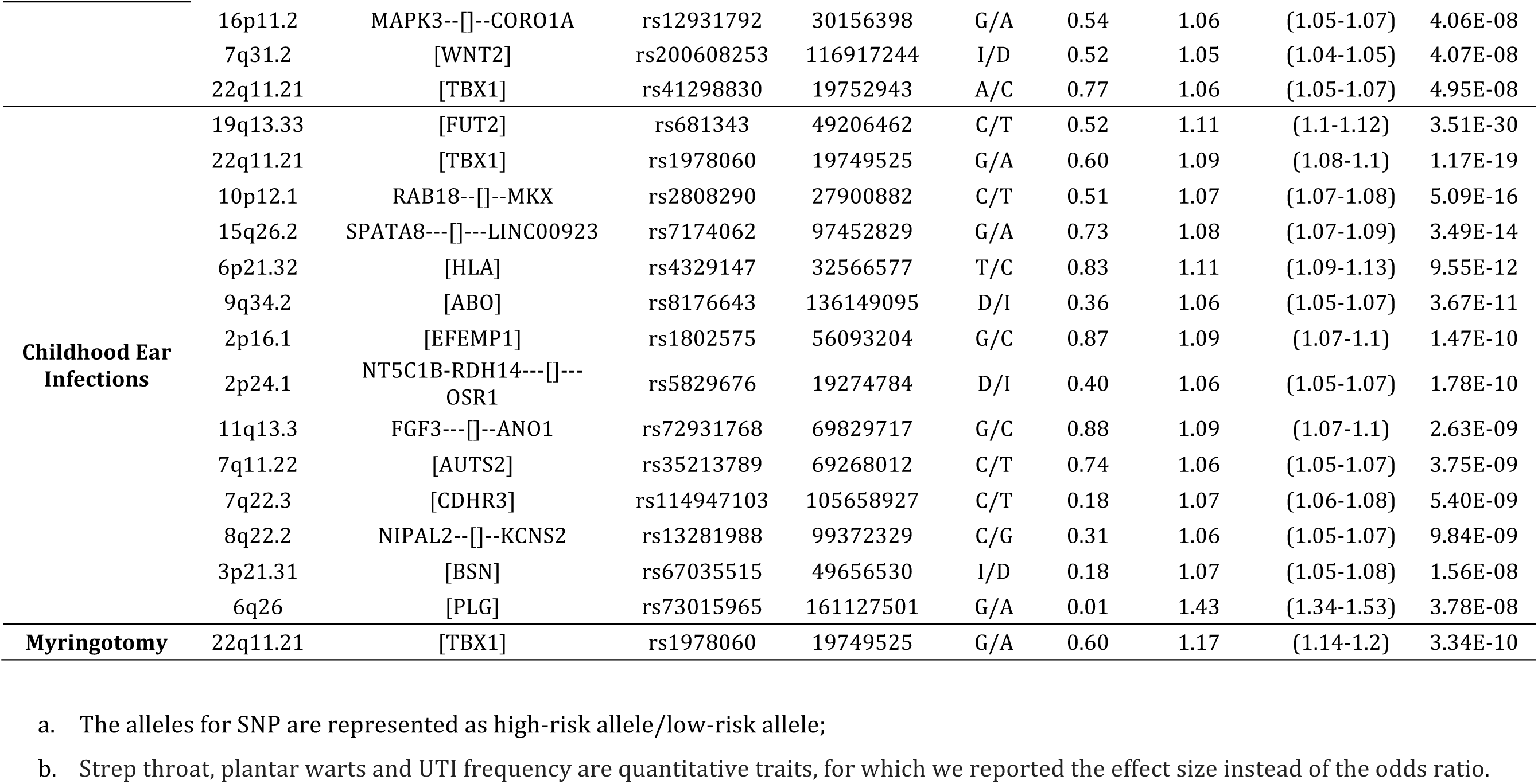
Genome-wide Significant Associations for Each Disease

**Figure 1.**
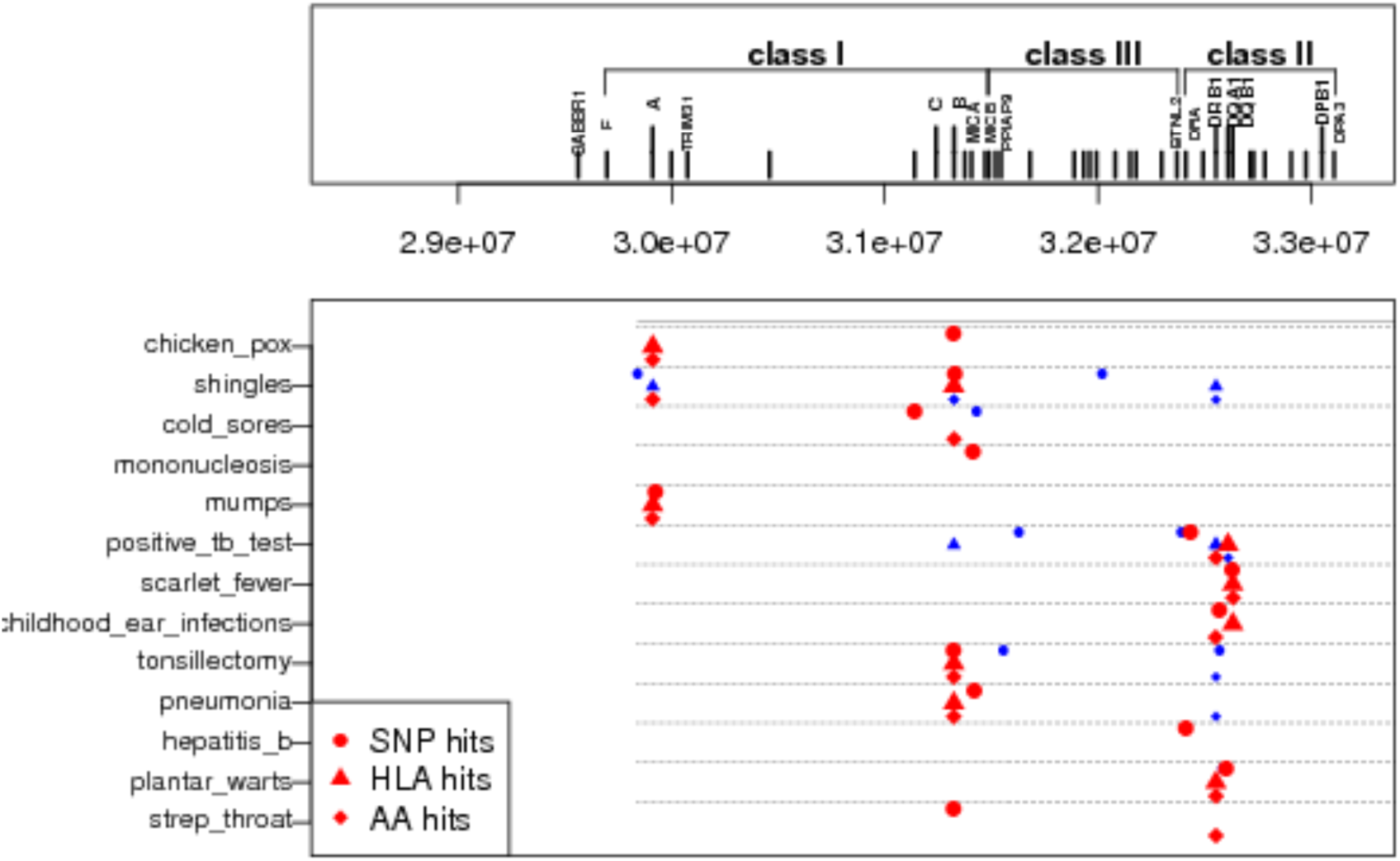
Summary of Independent HLA Signals. The strongest associated signals (red) and putative independent secondary signals (blue) are shown for each disease along with their location within the region. Independent secondary signals were defined as those with residual conditional association using the same significant threshold as the primary association signal mentioned in the main text.

**Table 3:**
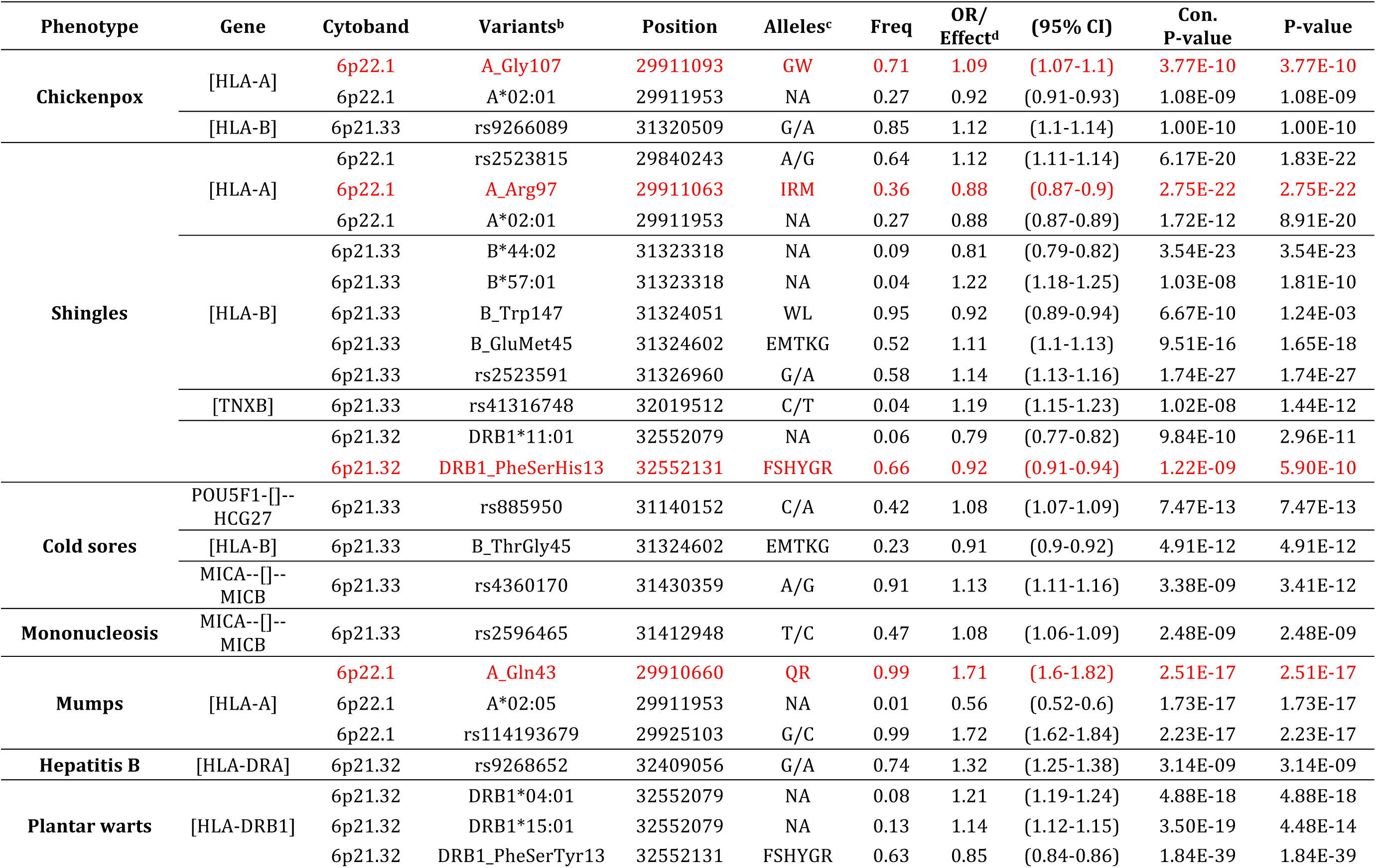
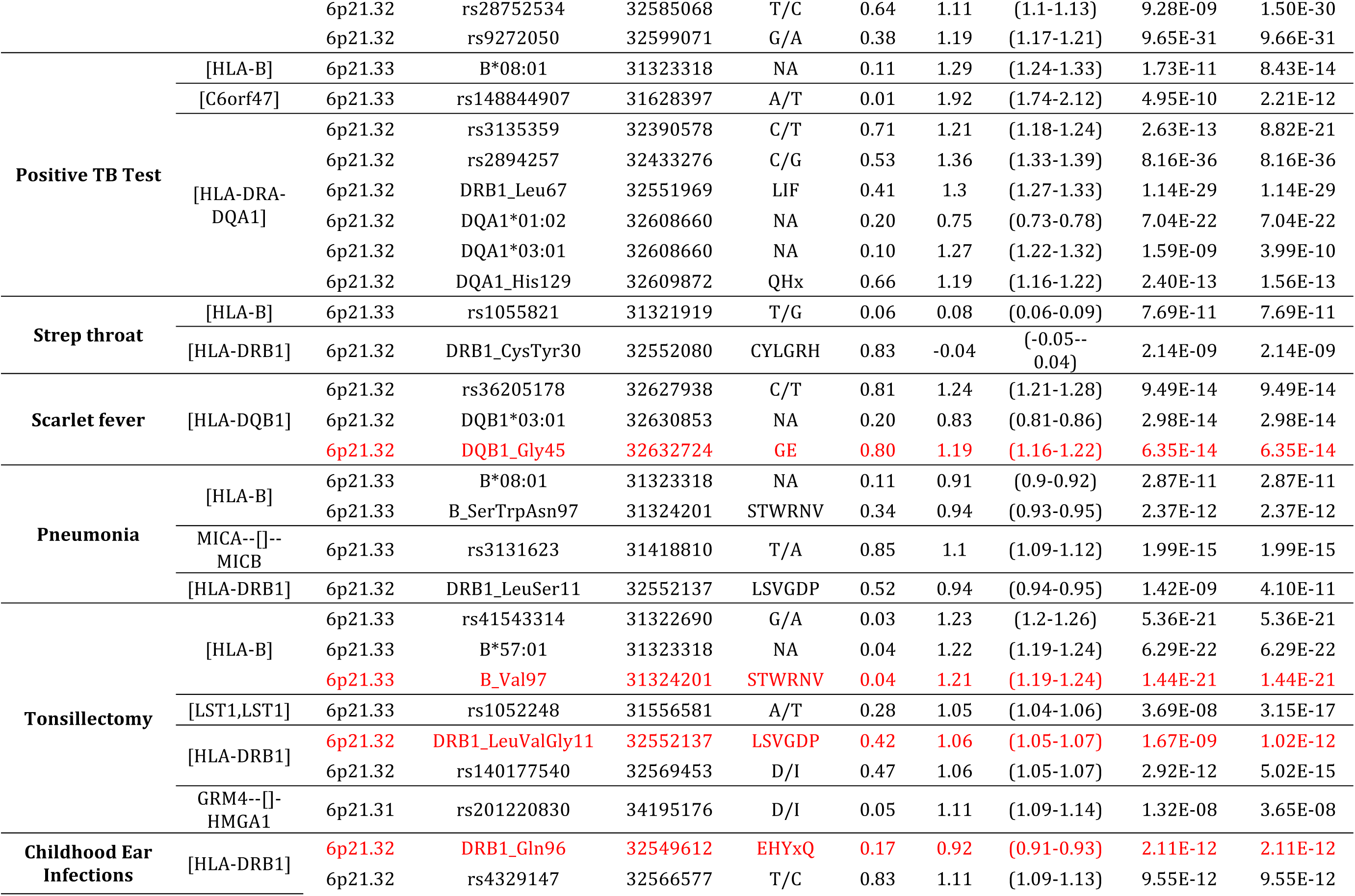
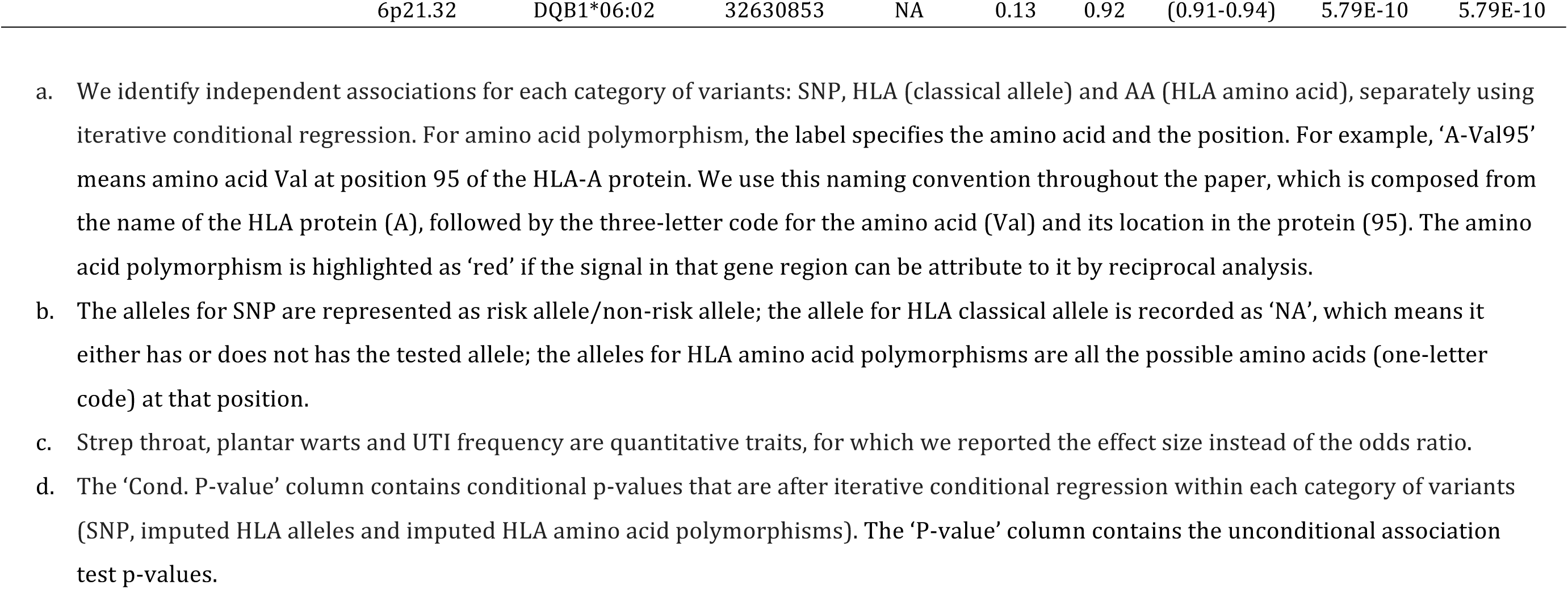
HLA Fine-mapping Results for Each Disease

## Genetic correlations

The pairwise LD score correlation on GWAS statistics (Figure 2, Supplementary Table 2) showed significant positive genetic correlations between strep throat and tonsillectomy (r_g_ = 0.85, se = 0.09, P = 1.89 × 10^−21^), between childhood ear infection and myringotomy (r_g_ = 0.88, se = 0.14, P = 1.27 × 10^−9^), and between chickenpox and shingles (r_g_ = 0.56, se = 0.16, P = 4 × 10^−4^), which are pairs of related phenotypes. A highly significant genetic correlation was found between tonsillectomy and childhood ear infections (r_g_ = 0.42, se = 0.05, P=8.19 × 10^−15^). Tonsillitis and ear infections are sometimes comorbid symptoms. A previous twin study suggested a substantial overlap in genetic factors influencing variations in liability to ear infection and tonsillitis^8^. In our GWASs, we identified the missense mutation (N397H) in *TBX1* as a significant susceptibility locus for both. Significant positive correlations were also observed for pneumonia with chronic sinus infections (rg = 0.72, se = 0.19, P = 1.0 × 10^−4^), childhood ear infections (rg = 0.39, se = 0.07, P = 1.47 × 10^−8^), colds last year (rg = 0.57, se = 0.12, P = 2.66 × 10^−6^) and strep throats (rg = 0.69, se = 0.13, P=1.0 × 10^−7^). These are all upper respiratory infections. The same viruses that cause colds and sore throat (if these infect the throat, sinuses and upper respiratory tract) can also cause pneumonia (if these reach the lungs). We also saw a high genetic correlation between urinary tract infection (UTI) and yeast infections (rg = 0.68, se = 0.08, P = 4.30 × 10^−16^). There was a phenotypic correlation (Pearson r = 0.41, P < 2.2 × 10^−16^) among women who reported both UTI and yeast infections in our cohort. These two infections can have similar symptoms and both cause discomfort in vaginal area, but different infectious agents cause them. The genetic correlation from LD score regression removes the correlation in GWAS summary statistics due to the correlation in phenotypes and thus suggests a shared host genetic susceptibility to UTI and yeast infections in females.

**Figure 2.**
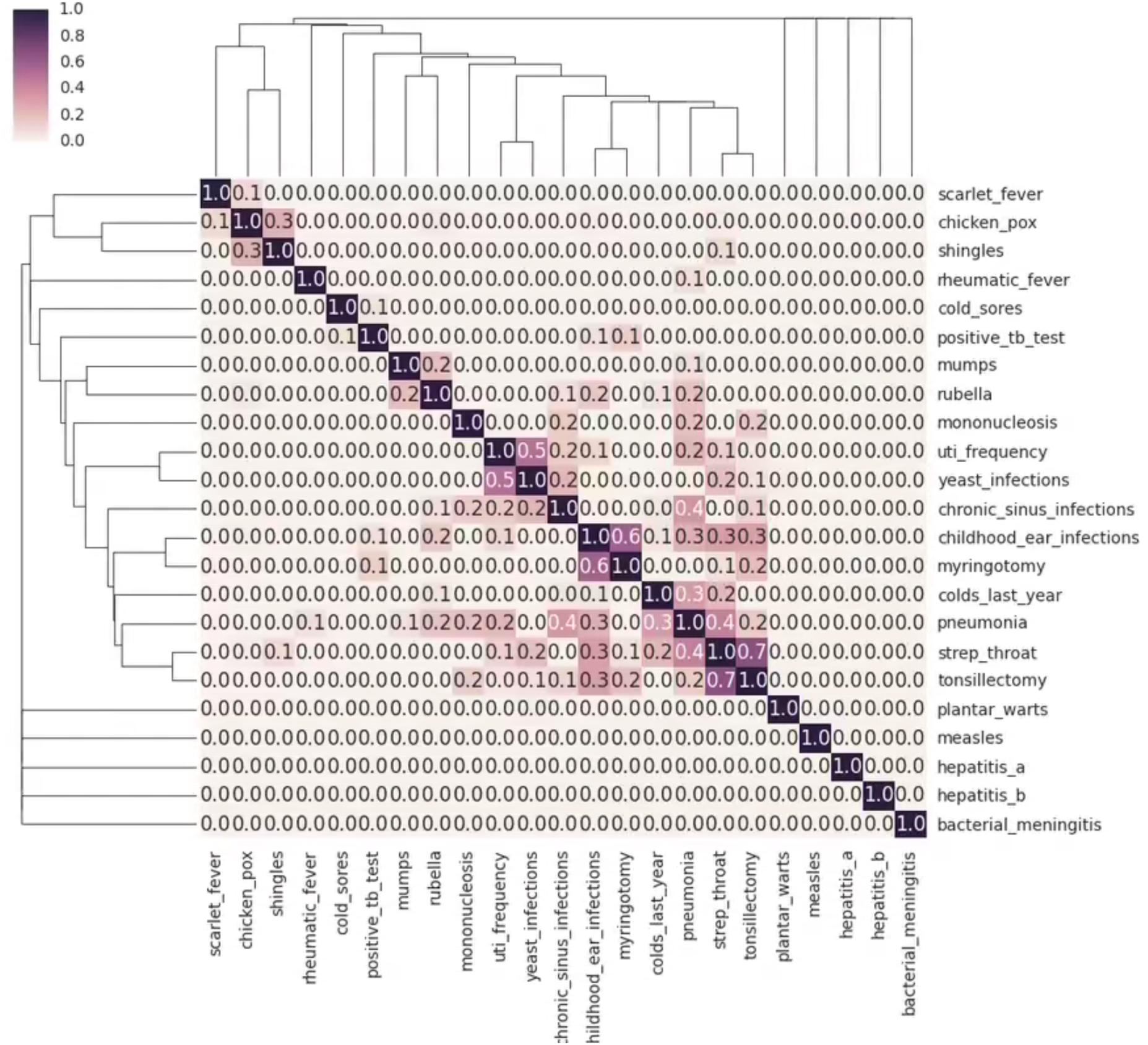
Heat map showing the genetic correlations. Each square [i, j] shows the lower bound of the 95% confidence interval of estimated genetic correlation (r_g_) between trait i and trait j, where i indexes row and j indexes columns. Darker colors represent larger genetic correlations. All negative values to zero. Phenotypes were clustered by ‘Nearest Point Algorithm’ implemented in python “Seaborn” package.

## Chickenpox (Varicella-zoster virus, Herpesviruses family)

Chickenpox, characterized by red, itchy bumps on the skin, is a highly contagious disease caused by primary infection with varicella-zoster virus (VZV). After the initial chickenpox infection, the virus remains dormant in the nervous system, though in approximately 20% of people it reactivates and manifests as shingles (see below). We identified independent associations in HLA-A and HLA–B in the class I region. On conditional analysis within HLA-A, the amino acid polymorphism HLA-A-Gly107 (P = 3.77 × 10^−10^) accounted for the HLA allele association (HLA-A*02:01, P = 1.08 × 10^−9^, conditional p-value is 0.90) in this interval. Although HLA-A-Gly107 is not located in the peptide-binding cleft, it is in high LD with HLA-A-Asp74, HLA-A-Gly62 and HLA-A-V95 (r^2^ > 0.95), which are all in the peptide-binding cleft. In HLA-B, only rs9266089 (P = 1.00 × 10^−10^) met the threshold for GWS.

## Shingles (Varicella-zoster virus, Herpesviruses family)

Shingles, also known as herpes zoster, is characterized by a painful band-shaped rash^9^ and is caused by reactivation of the VZV (see above). We identified multiple independent HLA signals for shingles. Within HLA-A, the amino acid polymorphism HLA-A-Arg97 (P = 2.75 × 10^−22^), located in the peptide-binding groove, accounted for most of the SNP effect (rs2523815, P = 1.83 × 10^−22^, conditional p-value is 0.04) and the HLA allele effect (HLA-A*02:01, P = 8.91 × 10^−20^, conditional p-value is 0.06). HLA-A*02:01 is implicated in both our chickenpox and shingles GWASs. The amino acid associations are different, but HLA-A-Arg97 is in high LD with HLA-A-Gly107 (r^2^ = 0.73). Within HLA-B, rs2523591 (P = 1.74 × 10^−27^) had a stronger association than HLA classical variants. There was a significant residual effect for rs2523591 after conditioning on HLA allele (conditional p-value is 1.71 × 10^−9^) or amino acid (conditional p-value is 3.13 × 10^−5^) associations in HLA-B region. The SNP rs2523591 is in LD with rs9266089 (r^2^ = 0.24, D’ = 0.98), which is a GWS signal in chickenpox GWAS, suggesting potential shared genetic factors for chickenpox and shingles. After adjusting for the signals in HLA-A and HLA-B regions, we observed independent secondary signals in the class III region (rs41316748, P=1.44 × 10^−12^) and in the class II region index by DRB1-PheSerHis13 (P = 5.9 × 10^−10^), which is in the peptide-binding cleft of HLA-DRB1.

We identified a variant upstream of *IFNA21* (rs7047299, P = 1.67 × 10^−8^) as a GWS association with shingles. None of the variants within 500kb and in moderate LD (r^2^ > 0.6) with the index SNP rs7047299 were coding, nor were they reported as expression quantitative trait loci (eQTL). *IFNA21,* a member of the alpha interferon gene cluster at band 9p21, encoded type I interferon and is mainly involved in innate immune response against viral infection. It have been shown to be involved in the pathogenesis of rubella^10^, and may also influence susceptibility to asthma and atopy^11^.

## Cold sores (type 1 herpes simplex virus, Herpesviruses family)

Over half of the US population suffers from cold sores (herpes simplex labialis), which are most commonly caused by the herpes simplex virus type I (HSV-1)^12^. Once someone has been infected, the virus usually cannot be eliminated and lies dormant in nerve cells where it may reactivate years later. Although most people are infected with HSV at some point in their lives, it is not clear why only some people suffer from cold sore outbreaks and reactivation^13^. We identified two independent GWS SNP associations in the HLA class I region. One is 2kb upstream of *POU5F1* indexed by rs885950 (P = 7.47 × 10^−13^) and the other is upstream of *HCP5* indexed by rs4360170 (P = 3.41 × 10^−9^). We also found a GWS association with HLA-B-ThrGly45 in the peptide-binding cleft of the HLA-B protein (P = 4.91 × 10^−12^). Upon conditioning on HLA-B-ThrGly45, the two SNPs had significant residual effects (conditional p-values are 6.28 × 10^−9^ and 1.32 × 10^−5^). However, the effect of HLA-B-ThrGly45 was largely removed (conditional p-value is 0.001) after conditioning on the two SNPs.

## Mononucleosis (Epstein-Barr virus, Herpesvirus family)

Over 90% of the world’s adult population is chronically infected with Epstein-Barr virus (EBV). Primary infection at childhood is usually asymptomatic or with only mild symptoms. Primary infection later in life is often accompanied by infectious mononucleosis (IM)^14^, commonly referred to as “mono” and characterized by fever, tonsillitis, and fatigue. A longitudinal study of IM in a cohort of 2,823,583 Danish children found evidence of familial aggregation of IM that warrants GWAS on IM disease etiology^14^. We identified the HLA class I region upstream of *HCP5* as a GWS association (indexed by rs2596465, P = 2.48 × 10^−9^). No GWS HLA allele or amino acid variant was identified.

## Mumps (mumps virus)

Mumps is an illness caused by the mumps virus. There is no disease modifying treatment for mumps infection. It is a rare disease in developed countries since the introduction of routine vaccination; however, outbreaks can still occur and mumps remains a significant health threat in developing countries^15^. We identified four independent GWS associations with mumps.

Variation in HLA-A, in the class I region, was significantly associated with mumps. The amino acid polymorphism HLA-A-Gln43 (P = 2.51 × 10^−17^) accounted for most of the SNP association (rs114193679, P = 2.23 × 10^−17^, conditional p-value is 0.19) and the HLA allele association (HLA-A*02:05, P = 1.73 × 10^−17^, conditional p-value is 0.26) in the HLA-A region.

We identified *FUT2* (indexed by rs516316, *P* = 9.63 × 10^−72^) and *ST3GAL4* (indexed by rs3862630, P = 1.21 × 10^−8^) as highly significant associations with mumps. Both belong to the glycosphingolipid (GSL) biosynthesis pathway, which was the most significant MAGENTA-analyzed pathway (Table 4: P = 1 × 10^−4^ and FDR = 0.003) identified using mumps GWAS data. This pathway also includes the *ABO* gene (indexed by rs8176643, P = 5.81 × 10^−5^). Glycosphingolipids (GSLs), found in the cell membranes of bacteria all the way to humans, play important biological roles in membrane structure, host-pathogen interactions, cell-cell recognition, and modulation of membrane protein function. Genetic variation in the human *FUT2* gene determines whether ABO blood group antigens are secreted into body fluids^16^. Detailed functional annotations of the genes identified in our mumps GWAS are provided in Supplementary table 3. The high-risk allele rs516316 (C) is in complete LD with rs601338 (A), a nonsense variant in *FUT2* that encodes the “non-secretor” (se) allele, which has been reported to provide resistance to Norovirus^17^ and susceptibility to Crohn’s disease^18,19^ and T1D^20^. The surface glycoprotein of mumps virus, hemagglutinin-neuraminidase, attaches to sialic acid receptors and promotes fusion and viral entry into host cells^21^. ABO antigens can modulate the interaction between pathogens and cell surface sialic acid receptors^22^; this presents a plausible mechanism by which secretor status could modify susceptibility to mumps infections. *ST3GAL4* encodes a member of the glycosyltransferase 29 family, a group of enzymes involved in protein glycosylation.

**Table 4:**
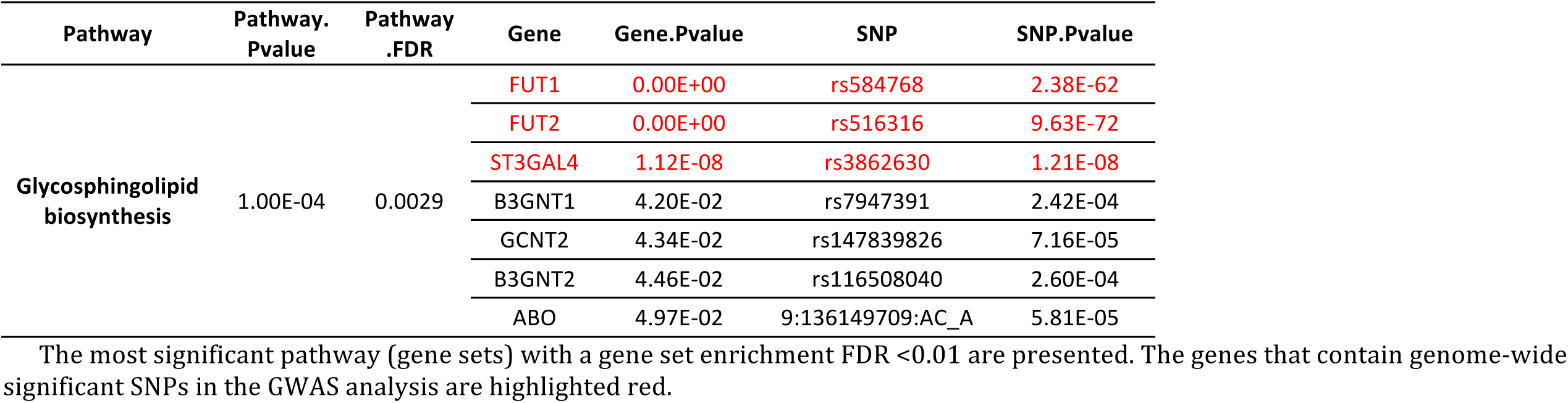
MEGANTA Pathway Analysis on Mumps GWAS

We also identified a significant association between mumps and a variant on chromosome 14 (rs11160318, *P* = 4.56 × 10^−12^) that is located in the intergenic region upstream of *BDKRB2.* The role of *BDKRB2* in mumps susceptibility is not clear, but the *BDKRB2* product has high affinity for intact kinins, which mediate a wide spectrum of biological effects including inflammation, pain, vasodilation, and smooth muscle contraction and relaxation^23^.

## Hepatitis B (hepatitis B virus)

Chronic hepatitis B virus (HBV) infection is a challenging global health problem. Several GWASs have been carried out to identify genetic loci involved in HBV susceptibility and immune response to hepatitis B vaccine, and the HLA-DRB1-HLA-DQB1 region has been consistently shown to be GWS^24–26^. In our GWAS of HBV infection we identified a GWS association with rs9268652 (P = 3.14 × 10^−9^, in HLA-DRA), consistent with the association of the HLA class II region with HBV susceptibility. Fine mapping analysis failed to identify GWS associations with HLA classical variants, but DQB1*06:02 was moderately associated with hepatitis B (P = 5.47 × 10^−6^). HLA-DQB1*06:02 is protective against HBV in our data, which is consistent with previous studies that the DRB1*15:01-DQB1*06:02 haplotype is associated with protection from type 1 autoimmune hepatitis and development of hepatocellular carcinoma^27^. Although chronic infection with the HBV has been linked to the development of hepatocellular carcinoma for more than 30 years^28^, the mechanism by which HBV infection leads to hepatocellular carcinoma is unclear.

## Hepatitis A (hepatitis A virus)

Infection with hepatitis A virus (HAV) can be asymptomatic or result in an acute illness characterized by flu-like symptoms and jaundice. Serious complications are rare and the hepatitis A vaccine is effective for prevention^29^. No large cohort study has been done for HAV infection. Our GWAS showed a suggestive association with *IFNL4,* indexed by rs66531907 (P = 5.7 × 10^−8^, OR = 1.23, 1kb upstream of *IFNL4). IFNL4* is located upstream of *IFNL3* (also known as *IL28B*). The high-risk allele rs66531907 (C) in our hepatitis A GWAS is in almost complete LD with rs8099917 (T) (r^2^ = 0.992), which has been associated with progression to chronic hepatitis C infection (HCV) and poor response to HCV therapy in multiple studies^30^.

## Plantar warts (human papillomavirus)

The human papillomavirus (HPV) causes plantar warts, which occur on the soles of the feet. Some strains of HPV are also associated with certain forms of cancer^2^. We identified two GWS associations with plantar wart. Our first association was with the HLA-DRB1 in class II region. The amino acid polymorphism DRB1_PheSerTyr13 (P = 1.84 × 10-39), in the peptide binding clefts of HLA-DRB1, was highly significant. Conditioning on it, the effects of HLA-DRB1*15:01 (P=4.48e-14, conditional p-value is 0.36), HLA-DRB1*04:01 (P = 4.88 × 10^−18^, conditional p-value is 0.001), and rs9272050 (P = 9.66 × 10^−31^, conditional p-value is 0.06) were largely removed, but a small residual effect remained for rs28752534 (P = 1.50 × 10^−30^, conditional p-value is 2.15 × 10^−8^).

We also identified a GWS association with rs6692209 (P = 5.25 × 10^−9^) near *LCE3E* in the Epidermal Differentiation Complex (EDC) on 1q21. The EDC comprises a remarkable density of gene families that determine differentiation of the human epidermis^31^. The high-risk allele rs6692209 (T) is in high LD (r^2^ = 0.86) with rs2105117 (A) (P = 2.98 × 10^−8^ in our GWAS), a missense variant in *LCE5A.* Variants in late cornified envelope (LCE) genes have been implicated in GWASs of atopic dermatitis (rs3126085-A, P = 6 x10^−12^, r^2^ = 0.047 with rs6692209)^32^ and psoriasis (rs4085613-A, P = 7 × 10^−30^, r^2^ = 0.002 with rs6692209)^33^, although the index SNPs identified in those GWASs were in low LD with our identified region. The skin is the primary interface with the external environment and provides a critical first barrier of the innate immune defense to infections^34^. While studies have linked the epidermal differentiation genes to various skin diseases, the role of the LCE proteins in epidermal biology has not been extensively studied. One study suggested that LCE proteins of groups 1, 2, 5 and 6 are involved in normal skin barrier function, while *LCE3* genes encode proteins that are involved in barrier repair after injury or inflammation^35^.

## Positive TB test (Mycobacterium tuberculosis)

Tuberculosis is a common and, in many cases, lethal infectious disease caused by various strains of mycobacteria, usually Mycobacterium tuberculosis. A chest x-ray and a sample of sputum are needed for a diagnosis of active TB disease^36^. The phenotype used in our GWAS was defined by asking customers whether they had had a positive response (red bumps) to a tuberculosis (TB) skin test. A positive TB test indicates that a person has at some time been exposed to and infected with TB bacteria. A negative TB test indicates that latent TB infection or TB disease is unlikely. In our GWAS of positive TB tests, the cases have either active TB disease, latent tuberculosis infections, or were vaccinated against TB, while controls may or may not have been exposed to TB bacteria and have never been infected.

We identified multiple independent HLA associations. In the most significant HLA-DRA-DQA1 region, conditioning on HLA-DRB1-Leu67 (P = 1.14 × 10^−29^) and HLA-DQA1-His129 (P = 1.56 × 10^−13^) removed the effect of DQA1*01:02 (P = 7.04 × 10^−22^, conditional p-value is 0.54), DQA1*03:01 (P = 3.99 × 10^−10^, conditional p-value is 0.36), and rs3135359 (P = 8.82 × 10^−21^, conditional p-value is 0.06); however, the effect of rs2894257 (P = 8.16 × 10^−36^, conditional p-value is 1.2 × 10^−7^) remained significant. Upon conditioning on the signals in the HLA-DRA-DQA1 region, secondary signals were detected in the class I (HLA-B*08:01, P=8.43 × 10^−14^) and class III regions (rs148844907 in C6orf47, P=2.21 × 10^−12^).

## Strep throat (group A streptococcus bacteria)

Strep throat (also called streptococcal tonsillitis or streptococcal pharyngitis) is characterized by fever, headache, severe sore throat, swollen tonsils and lymph nodes, and is caused by group A streptococcus (GAS) bacteria^37^. It is routinely treated with antibiotics. The symptoms of untreated strep throat typically resolve within a few days, but can sometimes cause complications such as rheumatic fever and kidney inflammation. We found a HLA-B class I region (index SNP rs1055821, P = 7.69 × 10^−11^) as a GWS association with strep throat. HLA-B*57:01 was almost GWS *(P* = 5.06 × 10^−8^). HLA-DRB1_CysTyr30 in class II region also showed a significant effect (P = 2.14 × 10^−9^). Conditioning on rs1055821, the effect of HLA-B*57:01 was removed (conditional p-value is 0.84) and the effect of HLA-DRB1_CysTyr30 was substantially decreased (conditional p-value is 0.005).

We also found an GWS association with a small insertion, rs35395352, in 1p36.23 (P = 3.90 × 10^−8^, effect = 0.03). None of the variants within 500kb and in moderate LD (r^2^ > 0.6) with this SNP were coding. However, two variants have been reported as cis-eQTLs: rs7548511 (P = 7.84 × 10^−8^, effect = 0.03, risk allele A, r^2^ = 0.96 with rs35395352) is an eQTL for *ENO1* (eQTL p-value is 8 × 10^−5^) in lymphoblastoid^38^ and rs11121129 (P = 9.62 × 10^−8^, effect = 0.03, risk allele G, r^2^ = 0.96 with rs35395352) is an eQTL for *TNFRSF9* (eQTL p-value is 4.4 × 10^−4^) in lymphoblastoid^39^. The product of ENO1, Alpha enolase, has been identified as an autoantigen in Hashimoto’s encephalopathy^40^ and a putative autoantigen in severe asthma^41^ and Behcet’s disease^42^. The *TNFRSF9-encoded* protein is a member of the TNF-receptor superfamily, which contributes to the clonal expansion, survival, and development of T cells. The same 1p36.23 region has been implicated in GWASs of psoriasis (rs11121129, risk allele A, P = 1.7 × 10^−8^, r^2^ = 0.96 with rs35395352)^43^. Interestingly, the strep throat risk allele rs11121129-A is protective against psoriasis.

Studies found that exacerbation of chronic psoriasis can be associated with streptococcal throat infections, and the T-cells generated by palatine tonsils can recognize skin keratin determinants in patients’ blood^44^.

## Scarlet fever (group A streptococcus bacteria)

Scarlet fever (also known as scarlatina) is caused by group A streptococcus (GAS) and is characterized by a bright red rash that covers most of the body, sore throat, and fever. It was a major cause of death before the discovery of effective antibiotics. We found a GWS association between scarlet fever and the HLA class II region. The amino acid polymorphism HLA-DQB1-Gly45 (P = 6.35 × 10^−14^), located in the peptide-binding cleft of HLA-DQB1, accounted for the SNP association (rs36205178, P = 9.49 × 10^−14^, conditional p-values is 0.83) and the HLA allele association (HLA-DQB1*03:01, P = 2.98 × 10^−14^, conditional p-values is 0.14) in this region. Although scarlet fever develops in some people with strep throat, we did not observe any cross disease effect between ‘scarlet fever’ and ‘strep throat’ in our analysis.

## Pneumonia (multiple origins)

Pneumonia is characterized by inflammation of the alveoli in the lungs and can be caused by a variety of organisms, including bacteria, viruses, and fungi. The most common cause of bacterial pneumonia in adults – isolated in nearly 50% of community-acquired cases-is Streptococcus pneumonia^45^. We found a GWAS association between pneumonia and a HLA class I region indexed by rs3131623 (P = 1.99 × 10^−15^). Conditioning on rs3131623, the residual effects of HLA-B-SerTrpAsn97 (P = 2.37 × 10^−12^, conditional p-value is 9.33 x10^−4^) and HLA-DRB1-LS11 (P = 4.10 × 10^−11^, conditional p-value is 3.89 × 10^−6^), both in the peptide-binding cleft, remained significant. There was also a residual effect left for rs3131623 (conditional p-value is 6.94 × 10^−5^) after conditioning on the two amino acid associations.

## Bacterial meningitis (Streptococcus pneumoniae, Group B Streptococcus bacteria and many other bacteria)

Bacterial meningitis is a severe infection causing substantial neurological morbidity and mortality worldwide. Streptococcus pneumoniae is the leading cause of bacterial meningitis and is associated with a 30% mortality rate^46^. We identified a GWS association with *CA10* (carbonic anhydrase X) indexed by rs1392935 (P = 1.19 × 10^−8^, intronic of CA10). None of the variants within 500kb and in moderate LD (r^2^ > 0.6) with rs1392935 were coding, nor were they reported as an eQTL. However, rs1392935 falls within an enhancer that is defined in the fetal brain. The protein encoded by *CA10* is an acatalytic member of the alpha-carbonic anhydrase subgroup and it is thought to play a role in the central nervous system, especially in brain development^47^.

## Yeast infection (Candida spp.)

A yeast infection, also called candidiasis, is usually a localized fungal infection of the skin or mucosal membranes, typically affecting the oral cavity, esophagus, gastrointestinal tract, urinary bladder, toenails, or genitalia^48^. Most yeast infections are caused by species of *Candida,* often *Candida albicans.* We found GWS associations between yeast infections and variants in *DSG1* (rs200520431, P = 1.87 × 10^−16^). The index SNP rs200520431 is intronic and in high LD (r^2^ > 0.8) with multiple missense mutations (rs8091003, rs8091117, rs16961689, rs61730306, rs34302455) in *DSG1.* The *DSG1* gene product is a calciumbinding transmembrane glycoprotein component of desmosomes in vertebrate epithelial cells. It connects the cell surface to the keratin cytoskeleton and plays a key role in maintaining epidermal integrity and barrier function^49^. This glycoprotein has been identified as the autoantigen of the autoimmune skin blistering disease pemphigus foliaceus^50^ and homozygous mutations in *DSG1* have been showed to result in severe dermatitis, multiple allergies, and metabolic wasting syndrome^49^. A variant downstream of *PRKCH* (indexed by rs2251260, P = 3.46 × 10^−10^) was also significantly associated with yeast infections. None of the variants within 500kb and in moderate LD (r^2^ > 0.6) with the index SNP rs2251260 were coding, nor were they reported as an eQTL. However, rs2251260 falls within strong enhancers defined in hepatocellular carcinoma. *PRKCH* is a member of a family of serine- and threonine-specific protein kinase and is predominantly expressed in epithelial tissues. The *PRKCH* protein kinase can regulate keratinocyte differentiation^51^. We also found a significant association with a variant in the 14q32.2 gene desert (rs7161578-T, p = 4.04 × 10^−8^). The index SNP is in high LD with many enhancer sequences defined in epidermal keratinocytes. A variant in the same region (rs7152623-A, p = 3 × 10^−15^, r^2^ = (+) 0.98 with rs7161578-T) was implicated in aortic stiffness^52^.

## Urinary Tract Infection (Escherichia coli, Staphylococcus saprophyticus, rarely viral or fungal)

About 80%-85% of urinary tract infections (UTIs) are caused by Escherichia coli (E. coli), and 5-10% are caused by Staphylococcus saprophyticus^53^. Very rarely, viruses or fungi may cause UTIs^54^. UTIs occur more commonly in women than men^55^ and in our dataset, significantly more female than male customers reported having two or more UTIs. We found an association between UTIs and variants in *PSCA* (indexed by rs2976388, *P* = 3.27 × 10^−10^). The index SNP rs2976388 falls within strong enhancers defined in epidermal keratinocytes. It is also in high LD (r2 = 0.95) with rs2294008 (P = 1.01 × 10^−9^), which is in the 5’-UTR promoter sequence of *PSCA* and can cause changes in transcriptional repressor CTCF motif. The PSCA gene variants conferred risk of UTI only in female when we further tested the effect in female and male cohort separately (P < 1e-11 in female, and P >0.1 in male). Prior GWASs have found associations between the *PSCA* gene and duodenal ulcer (rs2294008-C, p = 2e-33) and bladder cancer (rs2294008-T, p = 4 × 10^−11^)^56,57^. *PSCA* encodes a glycosylphosphatidylinositol-anchored cell membrane glycoprotein of unknown function. This glycoprotein was initially identified as a prostate-specific cell-surface marker^58^ and is overexpressed in a large proportion of prostate cancers, but also detected in bladder and pancreatic cancers^57^.

We also identified a variant intronic of *FRMD5* (rs146906133, P = 2.02 × 10^−8^) that is in LD with rs138763871 (r^2^ = 0.8, p = 1.75 × 10^−6^ in our GWAS), a missense variant in *STRC* (R1521W). The role of FRMD5 or STRC in UTIs is unclear.

## Tonsillectomy (multiple origins)

Tonsils are considered the first line of defense against respiratory infections. Previous twin studies have suggested a substantial genetic predisposition for recurrent tonsillitis^59^. Tonsillectomy is a surgery to remove the tonsils typically due to repeated occurrence of tonsillitis or tonsil hypertrophy. Our tonsillectomy GWAS revealed 35 independent chromosome regions reaching GWS. We observed strong HLA associations at HLA-B in the class I region. The amino acid polymorphism HLA-B-Val97 (P = 1.44 × 10^−21^), in the peptide-binding cleft of the HLA-B protein, accounted for most of the SNP effect (rs41543314, P = 5.36 × 10^−21^, conditional p value is 0.06) and HLA allele effect (HLA-B*57:01, P = 6.29 × 10^22^, conditional p value is 0.05) in this interval. We also noted additional independent signals in the class II and class III regions. Within HLA-DRB1 in the class II region, the effect of rs140177540 (P = 5.02 × 10^−15^, conditional p-value is 0.001) was largely removed after conditioning on DRB1_LeuValGly11 (P = 1.02 × 10^−12^) in the peptide-binding cleft of the HLA-DRB1 protein.

We detected 34 additional GWS associations (Table 2 and Supplementary Table 4), including signals from genes encoding tumor necrosis factors/receptors superfamily members *(LTBR, TNFRSF13B, TNFSF13B, CD40, TNFAIP3),* myotubularin (MTMR3), chemokines *(CXCL13),* mitogen-activated protein kinases/regulators *(IGFBP3, SPRED2, DUSP10, ST5, MAPK3),* SRC homology domain binding proteins *(SBK1, ST5, SH2B3),* disintegrins *(ADAM23, ADAMTS10),* various transcription factors *(NFKB1, HOXA2, FOXA1, IKZF1, MDFIC, TFEC, RERE, TBX1, FOXN1),* and signaling molecules *(WNT2, GNA12, ITSN1).* Many of these genes map to pathways involved in immune and inflammatory processes, while others are important regulators of embryonic development (e.g. most of the transcription factors), and a few are potentially involved in platelet production or hemostasis *(GNA12, SH2B3, RAP1GAP2, ADAM23, ADAMTS10).* When we tested the significant loci for enrichment in canonical pathways, the top over-represented pathway was ‘Intestinal immune network for IgA production’ (Table5, P = 2 × 10^−6^ and FDR = 3.5e-3). The genes that contributed significantly in this pathway include *LTBR* (rs10849448, P = 2.35 × 10^−35^), *TNFRSF13B* (rs34557412, missense variant, P = 2.66 × 10^−21^), *TNFSF13B* (rs200748895, P = 1.20 × 10^−17^) and *CD40* (rs6032664, P = 7.31 × 10^−12^). IgA is an antibody that plays a critical role in mucosal immunity. Tonsils belong to nasopharyngeal-associated lymphoid tissues and the generation of B cells is considered one of the major tonsillar functions; secretory dimeric IgA produced by the B cells is capable of preventing absorption and penetration of bacteria and/or viruses into the upper respiratory tract mucosa^60^. Tonsillectomy has been shown to significantly decrease levels of serum IgA and salivary secretory IgA levels^61^. Other identified significant signaling pathways include ‘TACI and BCMA stimulation of B cell immune system’ (P = 9 × 10^−4^ and FDR = 0.078), ‘ceramide signaling pathway’ (P = 3 × 10^−4^ and FDR = 0.08) and ‘Glucocorticoid receptor regulatory network’ (P = 5 × 10^−4^ and FDR = 0.099). The contributing genes are *TNFSF13B* (rs200748895, P = 1.2 × 10^−17^), *TNFRSF13B* (rs34557412, P = 2.66 × 10^−21^), *NFKB1* (rs23052, P = 4.54 × 10^−14^) and *MAPK3* (rs12931792, P = 4.06 × 10^−8^).

**Table 5:**
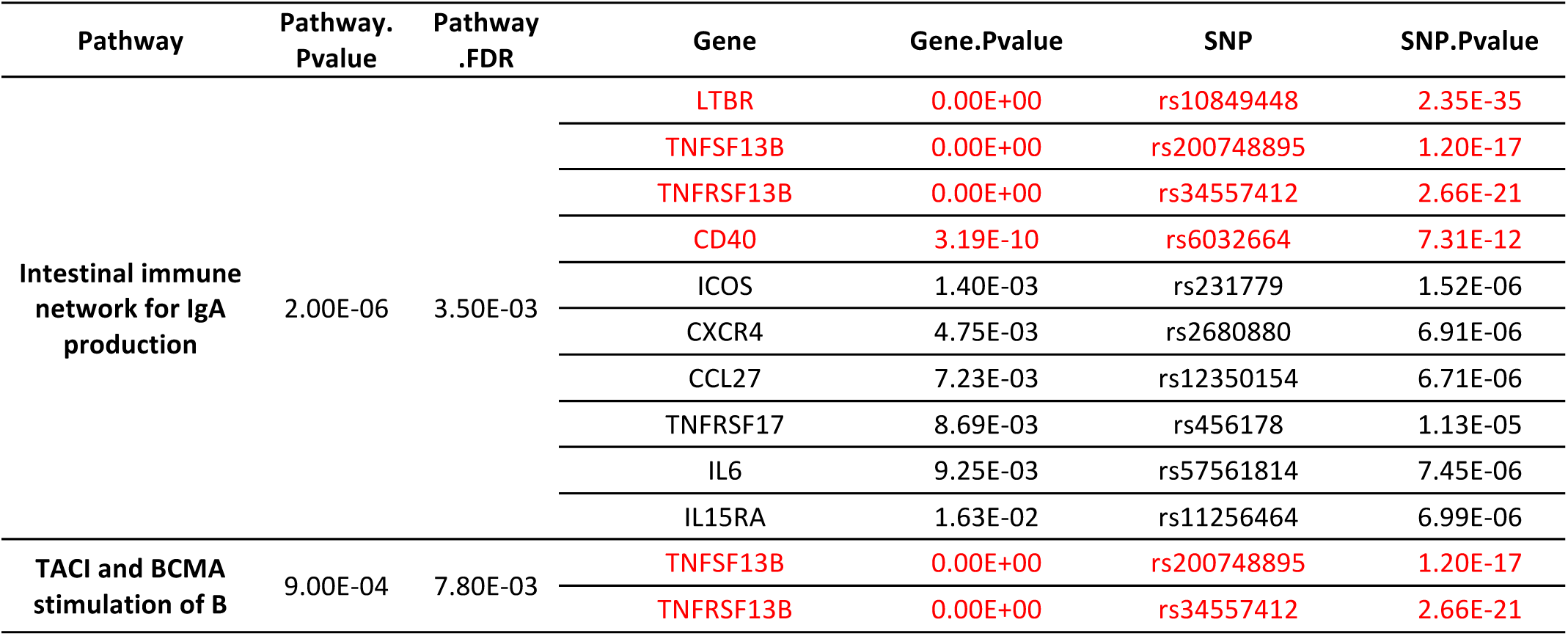
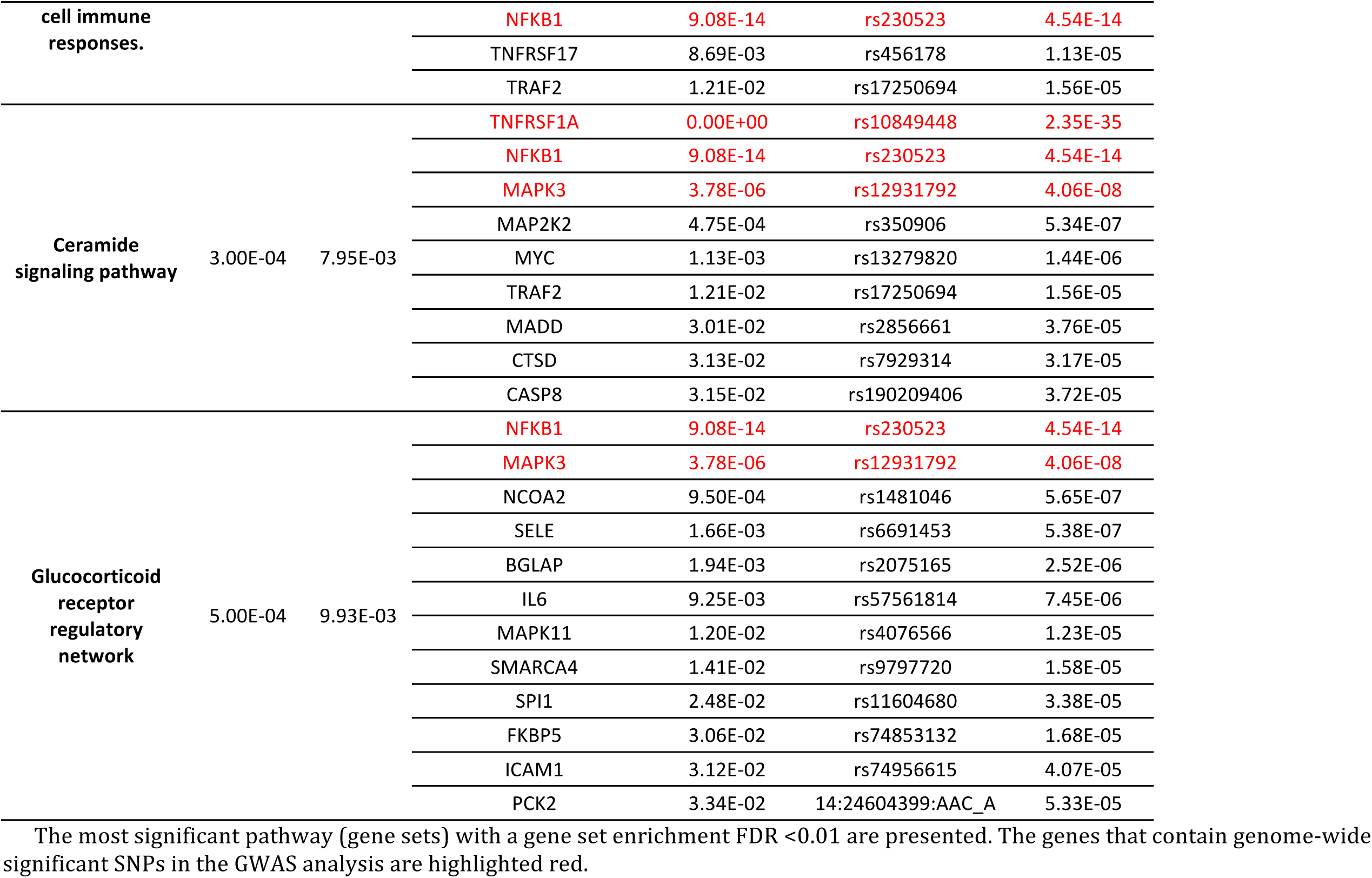
MEGANTA Pathway Analysis on Tonsillectomy GWAS

Although tonsillectomy has been performed for over 100 years, possible immunological effects of this procedure remain controversial. Many reports have demonstrated that in certain patients tonsillectomy is an effective therapy for psoriasis^44^ and rheumatoid arthritis (RA)^62^; the rationale for this effect is unknown. The ‘Intestinal immune network for IgA production’ pathway was also identified as the most significant pathway in a recent GWAS of IgAN^63^. In that study, they also linked the intestinal mucosal inflammatory disorders and inflammatory bowel disease (IBD) with risk of IgAN. Our tonsillectomy GWAS showed significant overlap of the identified loci with those autoimmune and inflammatory disorders^64^. We found five risk loci that are also associated with RA *(HLA, TNFIP3, CD40, SPRED2 and SH2B3);* four risk loci that are also associated with IBD *(HLA, MTMR3, TNFAIP3 and CD40)* and ulcerative colitis *(HLA, GNA12, MAPK3 and NFKB1);* three risk loci that are also associated with psoriasis *(HLA, SLC12AB and 1p36.23);* and two risk loci (*HLA* and *MTMR3*) that are shared with IgAN. We observed both concordant and opposing effects compared to these immune-mediated diseases. Our results may help to elucidate the connection between tonsillectomy and these diseases and may provide insight into clinical markers that could be used as indicators of tonsillectomy as a therapy for these diseases.

## Childhood ear infection (multiple origins)

Although ear infections can have a bacterial origin, they are often secondary to an upper respiratory infection such as a cold or sore throat, which can be caused by viruses or bacteria^65^. Children are more likely than adults to have ear infections because their Eustachian tubes are less effective at draining fluid and their immune systems are not fully developed. In our GWAS, we identified 14 regions that were significantly associated with childhood ear infection. Signals in the HLA region mapped to HLA-DRB1 in class II region. Our strongest association within HLA-DRB1 was with HLA-DRB1-Gln96 in the peptide-binding groove. After conditioning on HLA-DRB1-Gln96 (P = 2.11 × 10^−12^), the effects of rs4329147 (P = 9.55 × 10^−12^, conditional p value is 0.27) and DQB1*06:02 (P = 5.79 × 10^−10^, conditional p value is 0.85) were eliminated.

We found 13 additional GWS signals (Table 2 and Supplementary Table 5). Although MAGENTA pathway analysis did not identify significant pathways (Table 6), variants in *FUT2* and *ABO,* which were involved in the glycosphingolipid biosynthesis pathway and were implicated in our mumps GWAS, were also significantly associated with childhood ear infection. The risk allele rs681343(C) (*P* = 3.51 × 10^−30^) is a synonymous mutation in *FUT2* and is in almost complete LD with rs601338(G) (*r*^2^ = 0.9993), the ‘secretor’ (se) allele is associated with higher risk of childhood ear infection in our data. This is consistent with a previous report in which secretors were over-represented among patients with upper respiratory infections^66^. Our second most significant association was with a variant in *TBX1* (rs1978060, *P* =1.17 × 10^−19^). The low-risk allele rs1978060 (A) is in LD (r^2^ = 0.45) with rs72646967 (C), a missense (N397H) mutation in *TBX1* that was also implicated in our tonsillectomy GWAS. T-box genes encode transcription factors that play important roles in tissue and organ formation during embryonic development. In mice, *TBX1* haplo-insufficiency in the DiGeorge syndrome (22q11.2 deletion syndrome) region has been showed to disrupt the development of the pharyngeal arch arteries^67^ and the middle and outer ear^68,69^. We also identified other genes involved in developmental processes, including *MKX* (rs2808290, P = 5.09 × 10^−16^), *FGF3* (rs72931768, P = 2.63 × 10^−9^) and *AUTS2* (rs35213789, P = 3.75 × 10^−9^). The *MKX* product is an IRX family-related homeobox transcription factor and has been shown to play a critical role in tendon differentiation by regulating type I collagen production in tendon cells^70^. *FGF3* is a member of the fibroblast growth factor family, which is involved in a variety of biological processes including embryonic development, cell growth, morphogenesis, tissue repair and invasion. Study of similar genes in mouse and chicken suggested the role in inner ear formation^71^. *AUTS2* has been implicated in neurodevelopment and is a candidate gene for autism spectrum disorders and developmental delay^72,73^. Finally, we identified missense mutations in *CDHR3* (rs114947103, P = 5.40 × 10^−9^) and *PLG* (rs73015965, P = 3.78 × 10^−8^). The index SNP *rs114947103 is in almost complete LD with rs6967330 (r2=0.97), a missense variant in CDHR3, that* was recently identified in a GWAS as a susceptibility locus for asthma^74^. The biological role of *CDHR3* is not clear, but it is highly expressed in airway epithelium and belongs to the cadherin family that is involved in cell adhesion and epithelial polarity^74^. Mutations in the *PLG* gene could cause congenital plasminogen deficiency, which results in inflamed growths on the mucous membranes. Studies in mice have showed that *PLG* plays an essential role in protecting against the spontaneous development of chronic otitis media (middle ear infection) and have also suggested the possibility of using *PLG* for clinical therapy of certain types of otitis media^75^.

**Table 6:**
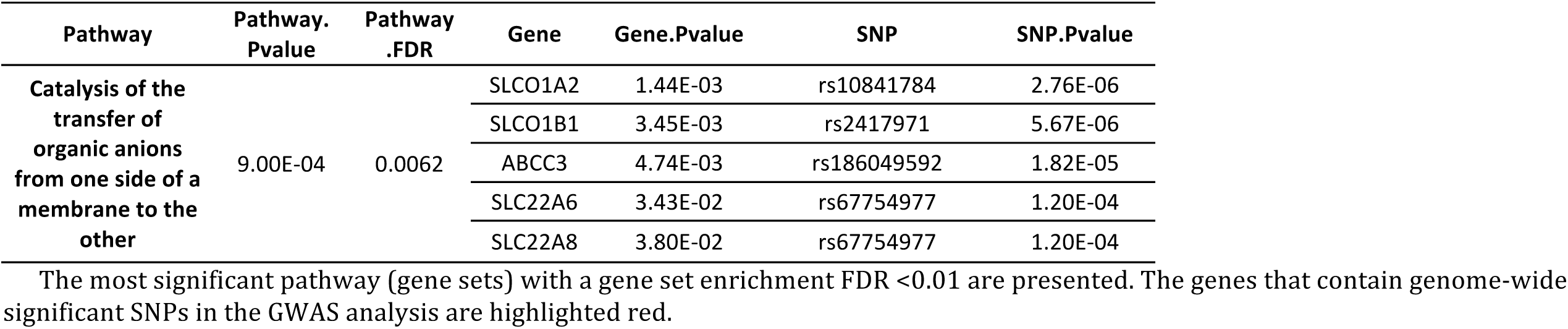
MEGANTA Pathway Analysis on Childhood Ear Infections GWAS

## Myringotomy (multiple causes)

Myringotomy is a surgical procedure in which a tiny incision is created in the eardrum to allow the release of middle-ear fluid. Before the invention of antibiotics, myringotomy without tube placement was a common treatment for severe or frequent acute otitis media^76^. The same variant in *TBX1* that was associated with tonsillectomy and childhood ear infection (see earlier) was also found to be significantly associated with myringotomy (rs1978060, *P* = 3.34 × 10^−10^).

## Discussion

We identified the greatest number of independent associations for tonsillectomy (n=35) and childhood ear infection (n=14), two relatively nonspecific phenotypes. For many of the infections, we only identified HLA associations. These findings may be due to our relatively larger sample sizes for the two traits and may also be due to the fact that tonsillectomy and childhood ear infections are heterogeneous phenotypes, influenced by a variety of pathogens and also by anatomical abnormalities. People undergo tonsillectomy for different reasons, including recurrent tonsillitis, obstructive sleep apnea, and nasal airway obstruction^77^. The causes of childhood ear infections are also diverse, involving different types of bacteria and viruses, and ear developmental defects^78^. Variants in the *TBX1* gene, which is essential for inner ear development^68,69^, were associated with both tonsillectomy and inner ear infections. Many genes that are involved in embryotic development were also identified in our tonsillectomy and childhood ear infection GWASs. The vulnerability to tonsillectomy and childhood ear infections may also be largely inherited. There were earlier studies calculated that heredity account for over 70% of susceptibility to recurrent ear infections in children^79,80^. Previous twin studies had suggested a substantial genetic predisposition for recurrent tonsillitis^59^ and sleep-disordered breathing^81^. According to the LD score regression on our GWAS data, tonsillectomy and childhood ear infections have relatively higher heritability on the observed scale than most of the other infections we studies. We did not calculate the liability scale heritability due to the lack of population prevalence information.

In the HLA region, we found that viral diseases — e.g., chicken pox, shingles, cold sores, mononucleosis (all caused by herpesvirus) as well as mumps (caused by mumps virus) — were mainly associated with variation in class I molecules (Figure 1). The bacterial diseases — specifically having a positive TB test (caused by Mycobacterium tuberculosis), scarlet fever (caused by GAS), and childhood ear infection (mostly caused by bacteria) — were mainly associated with variation in class II molecules. Tonsillectomy and pneumonia, caused by either bacteria or viruses, were associated with both class I and class II molecules. These observations are consistent with previous knowledge about antigen presentation; viruses mostly replicate within nucleated cells in the cytosol and produce endogenous antigens that are presented by class I MHC molecules, while bacteria grow extracellularly and are taken up by endosomal compartments where they are processed for presentation by class II MHC molecules^82^. These two intracellular pathways of protein processing may not be completely separate. Activation of CD8+, HLA class I-restricted T-cells by exogenous antigens has been reported^83^ and HLA class II-restricted CTLs that recognize endogenously synthesized antigen, such as HBV envelope antigens, have also been described^84,85^.

Some diseases, however, do not follow the common principles of antigen presentation. Plantar warts (caused by HPV) are different from other virus-causing diseases. HPV infections are exclusively intraepithelial and there is no viraemia or cytolysis in the infection cycle^86^. The primary response to HPV antigens is more likely to be initiated by the antigen-presenting cells (APCs) of squamous epithelia, the Langerhans cells (LCs). LCs capture antigens by macropinocytosis and receptor-mediated endocytosis, and then initiate class II processing of exogenous antigens^87^, which explains why our plantar warts GWAS mainly identified associations with MHC class II molecules. In other cases, our results suggest even more complex interactions. Strep throat (caused by GAS bacteria) has both MHC class I and class II associations. Some of our reported strep throat cases may represent secondary bacterial infections in response to a viral upper respiratory infection, meaning that the discovered MHC class I associations are actually with the virus infection that initiated the process^88^. Since most sore throats are caused by viruses^89^, another possibility is that some of the reported strep throat cases actually had a misdiagnosed or misreported viral infection. A final possibility is that HLA class I molecules can bind bacterial peptides derived from exogenous proteins that are internalized by endocytosis or phagocytosis, a process called cross-presentation^85,90^.

We failed to detect any GWS association for certain infectious diseases, including rubella, measles, chronic sinus infection, the common cold, or rheumatic fever. It is known that HLA affects responses to measles and rubella vaccinations^91,92^ and may thus also be associated with susceptibility. One limitation of our study that might affect findings with respect to measles and rubella is our lack of data on vaccination status. Since there is now a common vaccination for measles and rubella, misclassifying vaccinated people as controls would theoretically reduce our power to detect associations. This limitation might also affect the power of our shingles GWAS. The controls used for the shingles GWAS were not filtered for a positive history of chickenpox, though having chickenpox is a prerequisite for shingles. Another reason might be the heterogeneity of the phenotype. Colds are caused by several different viruses (mostly rhinoviruses and coronaviruses), and larger sample sizes may be required to identify alleles only associated with specific pathogens.

Although additional studies will be required to validate our associations, our findings are an important step towards dissecting the host genetic contribution to variation in response to infections. Research insights into infectious diseases will help to derive new diagnostic approaches and perhaps new therapies and preventions. This study may also help us to understand some of the autoimmune disorders that are associated with various infection triggers. Many of the identified associations were also found in autoimmune diseases. One postulated connection between infectious diseases and immunological disorders is that the immune cells that are activated in response to a pathogen epitope are also cross-reactive to self and lead to direct damage and further activation of other arms of the immune system^93^.

## Methods

### Subjects

We conducted GWASs of 23 infectious diseases. All participants were drawn from the customer base of 23andMe, Inc., a personal genetics company. All individuals included in the analyses provided informed consent and answered surveys online according to our human subjects protocol, which was reviewed and approved by Ethical & Independent Review Services, a private institutional review board (http://www.eandireview.com). All participants were of primarily European ancestry (>97% European ancestry) as determined through an analysis of local ancestry^94^. A maximum set of unrelated individuals was chosen using a segmental identity-by-descent (IBD) estimation algorithm^95^. Individuals were defined as related if they shared more than 700 cM of IBD, including regions where the two individuals share either one or both genomic segments IBD. This level of relatedness (involving ~20% of the human genome) corresponds approximately to the minimal expected sharing between first cousins in an outbred population.

Table 1 shows our discovery sample sizes, drawn from more than 650,000 genotyped customers who reported via web-based questionnaires whether they had been diagnosed with one or more infectious diseases. (For more detail on survey questions used to collect the phenotype data, see the Supplementary Notes).

### Genotyping and SNP imputation

DNA extraction and genotyping were performed on saliva samples by National Genetics Institute (NGI), a CLIA-certified laboratory that is a subsidiary of Laboratory Corporation of America. Samples were genotyped on one of four genotyping platforms. The V1 and V2 platforms were variants of the Illumina HumanHap550+ BeadChip and contained a total of about 560,000 SNPs, including about 25,000 custom SNPs selected by 23andMe. The V3 platform was based on the Illumina OmniExpress+ BeadChip and contained a total of about 950,000 SNPs and custom content to improve the overlap with our V2 array. The V4 platform in current use is a fully custom array of about 950,000 SNPs and includes a lower redundancy subset of V2 and V3 SNPs with additional coverage of lower-frequency coding variation. Samples that failed to reach 98.5% call rate were re-analyzed. Individuals whose analyses failed repeatedly were re-contacted by 23andMe customer service to provide additional samples, as is done for all 23andMe customers.

Participant genotype data were imputed using the March 2012 “v3” release of the 1000 Genomes Project reference haplotypes^96^. We phased and imputed data for each genotyping platform separately. First, we used BEAGLE^97^ (version 3.3.1) to phase batches of 8,000 to individuals across chromosomal segments of no more than 10,000 genotyped SNPs, with overlaps of 200 SNPs. We excluded SNPs with minor allele frequency (MAF) < 0.001, Hardy-Weinberg equilibrium *P* < 1 × 10^−20^, call rate < 95%, or large allele frequency discrepancies compared to the European 1000 Genomes Project reference data. Frequency discrepancies were identified by computing a 2×2 table of allele counts for European 1000 Genomes samples and 2000 randomly sampled 23andMe customers with European ancestry, and identifying SNPs with a chi squared P < 10^−15^. We imputed each phased segment against all-ethnicity 1000 Genomes haplotypes (excluding monomorphic and singleton sites) using Minimac2^98^ with five rounds and 200 states for parameter estimation.

For the non-pseudoautosomal region of the × chromosome, males and females were phased together in segments, treating the males as already phased; the pseudoautosomal regions were phased separately. We then imputed males and females together using Minimac as with the autosomes, treating males as homozygous pseudo-diploids for the non-pseudoautosomal region.

### GWAS analysis

For case control comparisons, we tested for association using logistic regression, assuming additive allelic effects. For quantitative traits, association tests were performed using linear regression. For tests using imputed data, we use the imputed dosages rather than best-guess genotypes. We included covariates for age, gender, and the top five principal components to account for residual population structure. The association test p-value was computed using a likelihood ratio test, which in our experience is better behaved than a Wald test on the regression coefficient. Results for the × chromosome were computed similarly, with men coded as if they were homozygous diploid for the observed allele.

### Genetic correlations

We computed LD scores as previously described^99^ using the samples in the 1000 Genomes Project reference panel (phase 3 verion 5a) with a MAF cutoff of 5%. We calculated the pairwise genetic correlation (r_g_) on the GWAS summary statistics (including MHC region) using LD score regression^99^. The cluster map of genetic correlation, base on the lower bound of the 95% confident interval of calculated r_g_, was plotted using ‘Nearest Point Algorithm’ implemented in python “Searborn” package (https://stanford.edu/~mwaskom/software/seaborn/index.html). We found no significant negative correlations in our data, thus we set all negative values to zero.

### SNP Function Annotation

To explore whether any of the significant SNPs identified might link to functional mutations or have potential regulatory functions, we used the online tools HaploReg (http://www.broadinstitute.org/mammals/haploreg/haploreg.php) to confirm the location of each SNP in relation to annotated protein-coding genes and/or non-coding regulatory elements. We queried only the variants that were within 500kb of and in moderate LD (r^2^>0.6) with the index SNP.

### Imputation of HLA classical alleles and respective amino acid variations

HLA imputation was performed with HIBAG^100^, an attribute bagging based statistical method that comes as a freely available R package and includes a pre-fit classifier. This classifier was trained from a large database of individuals with known HLA alleles and SNP variation within the HLA region. Over 98% of the tagging SNPs used in HIBAG were genotyped and passed quality control on 23andMe’s platform. We imputed allelic dosage for HLA-A, B, C, DPB1, DQA1, DQB1, and DRB1 loci at four-digit resolution. We used the default settings of HIBAG and the run time for 100,000 samples was about 10 hours on our cluster.

Using an approach suggested by P. de Bakker^101^, we downloaded the files that map HLA alleles to amino acid sequences from https://www.broadinstitute.org/mpg/snp2hla/ and mapped our imputed HLA alleles at four-digit resolution to the corresponding amino acid sequences; in this way we translated the imputed HLA allelic dosages directly to amino acid dosages. We encoded all amino acid variants in the 23andMe European samples as biallelic markers, which facilitated downstream analysis. For example, position 45 of HLA-B protein had five different alleles (E: Glu, M: Met, T: Thr, K: Lys, G: Gly), we first encoded the position using five binary markers, each corresponding to the presence or absence of each allele (e.g. HLA-B-Gly45 tags Gly at position 45 of HLA-B protein). For positions having three or more alleles, we also created markers that tag multiple alleles, each corresponding to the presence or absence of the multiple alleles (e.g. HLA-B-ThrGly45 tags Thr and Gly at position 45 of HLA-B protein). Thus, we created binary indicators for all possible combinations of amino acid variants. We use this naming convention for amino acid polymorphisms throughout this paper.

We imputed 292 classical HLA alleles at four-digit resolution and 2395 bi-allelic amino acid polymorphisms. Similar to the SNP imputation, we measured imputation quality using *r^2^,* which is the ratio of the empirically observed variance of the allele dosage to the expected variance assuming Hardy-Weinberg equilibrium. The imputation quality (*r*^2^) of the top associated HLA alleles and amino acids are in Supplementary Table 6.

### HLA region fine mapping

To test associations between imputed HLA allele/amino acid dosages and phenotypes, we performed logistic or linear regression using the same set of covariates used in the SNP-based GWAS for that phenotype. We applied a forward stepwise strategy, within each type of variant, to establish statistically independent signals in the HLA region. Within each variant type(e.g. SNP, HLA allele, and HLA amino acid), we first identified the most strongly associated signals (lowest p-value) for each disease and performed forward iterative conditional regression to identify other independent signals if the conditional p-value was < 5 × 10^−8^. All analyses controlled for sex and five principal components of genetic ancestry. The p-values were calculated using a likelihood ratio test. The iterative conditional regression dissected HLA signal into independent HLA associations. Within each identified HLA locus (HLA-A, B, C, DPB1, DQA1, DQB1, and DRB1), We further carried out reciprocal analyses, which are the conditional analyses across variants types, to see if the association can be attributed to the amino acid polymorphism within each HLA locus.

### Pathway analysis

In order to better understand how multiple genes in the same pathway may contribute to certain infections, we performed pathway analysis using Meta-Analysis Gene-set Enrichment of variaNT Associations (MAGENTA)^102^, which tests for enrichment of genetic associations in predefined biological processes or sets of functionally related genes, using GWAS results as input. We used gene sets of 1320 canonical pathways from the Molecular Signatures Database (MsigDB) compiled by domain experts^103^ and default settings of MAGENTA tool.

## Acknowledgements

We thank the customers of 23andMe for participating in this research and the employees of 23andMe for their contributions to this work.

